# Severe deviation in protein fold prediction by advanced AI: a case study

**DOI:** 10.1101/2024.08.20.608820

**Authors:** Jacinto López-Sagaseta, Alejandro Urdiciain

## Abstract

Artificial intelligence (AI) and deep learning are making groundbreaking strides in protein structure prediction. AlphaFold is remarkable in this arena for its outstanding accuracy in modelling proteins fold based solely on their amino acid sequences. In spite of these remarkable advances, experimental structure determination remains critical. Here we report severe deviations (>30 Å) between the experimental structure of a two-domain protein and its equivalent AI-prediction. These observations are particularly relevant to the relative orientation of the domains within the global protein scaffold. Significant divergence between experimental structures and AI-predicted models echoes the presence of unusual conformations, insufficient training data and high complexity in protein folding that can ultimately lead to current limitations in protein structure prediction.

## Main Text

The potential for determining protein structures based solely on the amino acidic sequence is rapidly evolving, driven primarily by the availability of vast numbers of experimental structures that artificial intelligence and deep learning resources can exploit. Most recently, AlphaFold^1^ has led transformative effects in the field of structural biology by enabling fast and unprecedented accurate prediction of protein structures. Despite these highly significant contributions, several limitations and challenges^2,3^ in protein structure determination cannot yet be addressed by computational procedures.

Recently^4^, we have described the experimental structure of a marine sponge receptor via X-ray crystallography with a resolution of 1.6 Å **(Table 1)**. This fragment corresponds to two tandem Ig-like domains that are part of the extracelullar region of this receptor known as SAML (Sponge Adhesion Molecule, long form). Given the lack of a good homologue that could be used to solve the structure through molecular replacement, we used the predicted AlphaFold equivalent. This attempt failed to yield a structure solution. However, an alternative trial using, separately, the individual Ig domains led to the solution of the target structure using the same procedure^5^. This fact prompted us to interrogate the accuracy of the AlphaFold predicted model (AF-Q9U965-F1). Thus, we confronted it with the atomic coordinates obtained with the experimental structure. Protein-protein alignments and superpositions through the C-alpha atoms readily showed remarkable discrepancies (RMSD 7.735 Å) between the predicted and experimental structures **(Fig. S1)**.

**Table 1.** Diffraction data collection and refinement statistics.

|  | SAML (PDB 8OVQ) |
| --- | --- |
| Resolution range (Å) | 77.8- 1.6 (1.64 - 1.61) |
| Space group | P12<sub>1</sub>1 |
| Unit cell | 33.08 41.16 77.80 90 90 90 |
| Total reflections | 160776 (15963) |
| Unique reflections | 27240 (2722) |
| Redundancy | 4.1 (3.7) |
| Completeness (%) | 99.40 (99.82) |
| Mean I/sigma(I) | 9.3 (1.5) |
| Wilson B-factor | 20.56 |
| R-merge | 0.067 (0.74) |
| CC<sub>1/2</sub> | 0.997 (0.576) |
| Reflections used in refinement | 27228 (2722) |
| Reflections used for R-free | 4063 (414) |
| R-work | 0.169 (0.304) |
| R-free | 0.216 (0.336) |
| RMS (bonds) | 0.012 |
| RMS (angles) | 1.85 |
| Ramachandran favored (%) | 98 |
| Ramachandran allowed (%) | 2.0 |
| Ramachandran outliers (%) | 0.0 |
| Average B-factor | 32.5 |

Statistics for the highest-resolution shell are shown in parentheses.

Yet, more striking differences were noticed when the structures were compared by aligning them through either the N- or the C-terminal Ig domains. In both cases, the results show evident architecture mismatches. The orientation of the free Ig-domains relative to the aligned Ig-domains shows a strong deviation in the predicted models **(Fig. 1)**. These discrepancies are particularly remarkable as we would expect the alignments of the N- and C-terminal Ig-domains to be reasonably close to the experimental structure, regardless of the aligned reference region, whether N- or C-terminal. Thus, these observations indicate that the prediction of the interdomain spatial relationship does not match that of the experimental structure. These findings would explain why our initials attempts failed to find a plausible solution in the molecular replacement procedure. On the contrary, the alignment of the individual domains yielded closely related structures **(Fig. S2)**, as indicated by the root mean square deviations (RMSD) values below 0.9 Å (**Figs. S2A and S2B)**.

**Fig 1.**
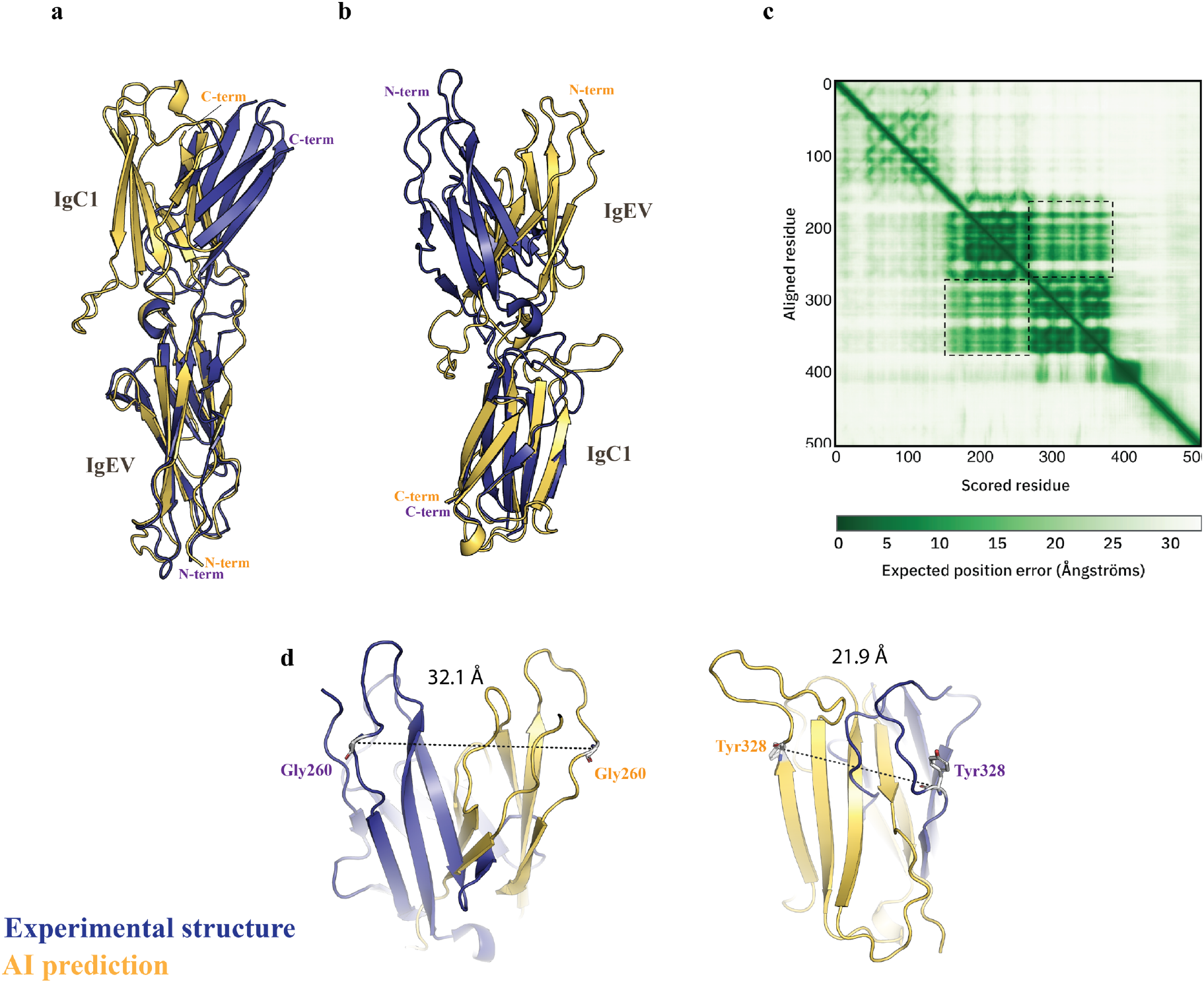
Fold deviation in the predicted AI-model. **A-B**, the experimental structure (deepblue) and AI-derived model (yelloworange) are represented in cartoon mode and are superposed through the alpha carbons of the N-terminal (a) or C-terminal (b) Ig-domains. The N-terminal early-variable Ig-like (EV) domain and the C-terminal Ig-like C1^4^-set are indicated. **C**, graphic plot (source AlphaFold) accounting for the PAE error of the predicted AlphaFold model. The dashed squares indicate the areas for expected errors of the Ig interdomain spatial relationships. **D**, the position of the residues showing the strongest deviations between the X-ray structure and the AlphaFold prediction are displayed in Å.

Further evident discrepancies can be unambiguously noticed at a more local level. While these are mainly found in the loop regions, the impact of these discrepancies leads to additional deviations in the location and orientation of some beta strands that are part of the rigid core of the Ig-like β sandwich. This is expected as loops are usually associated with high flexibility and is also consistent with the AlphaFold anticipation of expected low accuracy for these regions. Nevertheless, discrepancies hit not only from a folding perspective, but concerning the type of secondary structure. For instance, and according to the Dictionary of Secondary Structure in Proteins (DSSP) for secondary structure assignment^6^, the A beta strand of the C-terminal Ig-domain expands from Leu274 to Leu283 (**<u>LIVEVDSSGL</u>**) in the X-ray structure, while only the portion defined by residues Ile275 to Asp279 (L**<u>IVEVD</u>**SSGL) is considered as a β strand motif by the AI model **(Fig. S3A)**. Moreover, an additional region with a higher confidence level (90 > pLDDT > 70) presents also an unmatched secondary structure assignment **(Fig. S3B)**. Here, the model prediction assigns a beta strand secondary motif (**FNITPRY**), while the X-ray structure indicates that only a fragment of this sequence corresponds to a beta strand **(Fig. S3B)**, with the remaining fraction showing a loose area with no secondary structure (F**NITP**RY).

Yet, and beyond the local discrepancies that can be foreseen by the predictor confidence metrics, what is particularly relevant in this case study is the aforementioned evident divergence that exists in the spatial relationship between the two Ig-domains that conform the target protein. Two major factors can contribute to these deviations. On the one hand, in the X-ray structure, the protein surrounding moiety (buffer, ions and other small molecules), the crystal packing contacts and other potential environmental factors might bias a particular conformation in the experimental structure. On the other hand, current AI limitations might not optimally address interdomain interactions, leading to inaccurate predictions in multidomain proteins.

AlphaFold outputs are accompanied by thorough details describing the confidence metrics for a particular target. These are based on a predicted local-distance difference test (pLDDT), a predicted aligned error (PAE), estimates of the predicted template modeling (pTM) and the interface predicted template modeling (ipTM) scores^1^. Thus, any user is informed about the accuracy of the predicted model. The PAE plot calculated by AlphaFold accounts for the accuracy of the position and orientation of two independent domains relative to each other within a single protein. In this case, the PAE plot suggests moderate to low error values, which would anticipate, relatively, a modest error in the expected position of these domains relative to each other **(Fig. 1c)**. However, the structural comparison indicates a strong disagreement for the relative position of the N-terminal and C-terminal domains **(Figs. 1A-B and 1D)**.

The observed differences in the orientation of the single domains relative to each other between the predicted and the experimental structures highlight the complexity of these predictions, as mentioned, especially in multidomain proteins. A precise definition of the folding mode in multidomain proteins is pivotal for the advance of biomedicine: their role in protein-protein interactions, the potential for allosteric modulation, or their utility in structure-based drug design are just some examples. It is expected that the continuous expansion of experimental structures and subsequent training models will lead to improved algorithms that account for high-accuracy inter-domain orientation.

## Methodology

All the methodology is described in the preprint by Urdiciain *et al*.: Unusual traits shape the architecture of the Ig ancestor molecule. doi: https://doi.org/10.1101/2024.07.22.604567.

## Acknowledgements

We thank the staff of the Xaloc beamline at ALBA Synchrotron for their assistance with X-ray diffraction data collection.

## Funding

Ramón y Cajal, Grant RYC-2017-21683, Ministry of Science and Innovation, Government of Spain (JLS). Alejandro Urdiciain is a recipient of a Margarita Salas contract funded by UPNA and the Ministry of Universities of Spain within the Plan of Recovery, Transformation and Resilience and the European Recovery Instrument Next Generation EU.

## Author contributions

Conceived research project: JLS; Performed experiments: AU, JLS; Data analysis: JLS, AU; Draft writing: JLS.

## Competing interests

The authors declare that they have no conflict of interest.

## Data availability

Atomic coordinates and structure factors for SAML is available in the Protein Data Bank under the accession codes 8OVQ.

**Fig S1.**
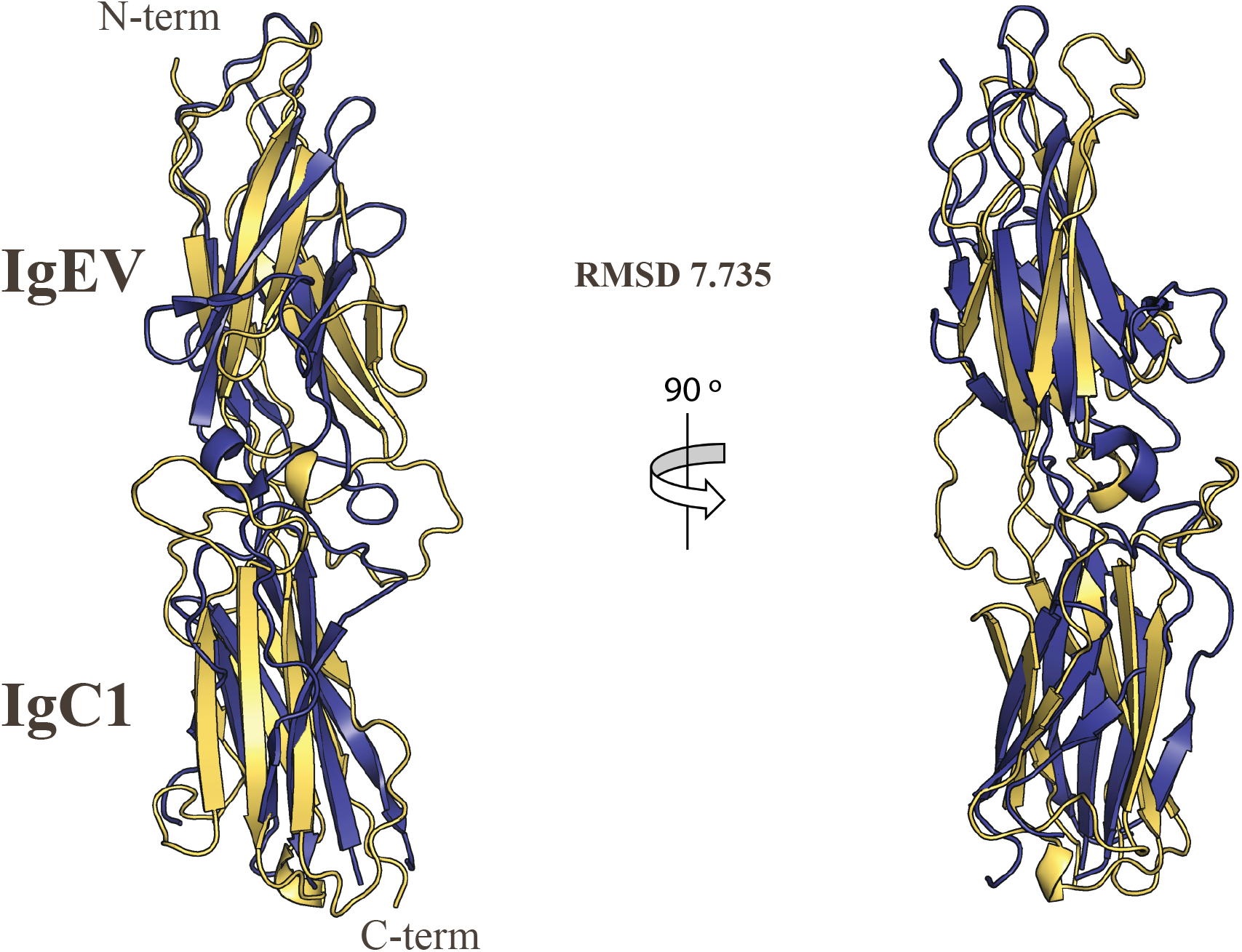
Structural alignment of the X-ray SAML structure and the predicted AlphaFold model. Both structures are displayed as ribbons colored in deepblue (X-ray structure) and yelloworange (AlphaFold model) and superposed through the alpha carbons. A second view is shown at right with both structures rotated 90 º with respect to the original view (left panel).

**Fig S2.**
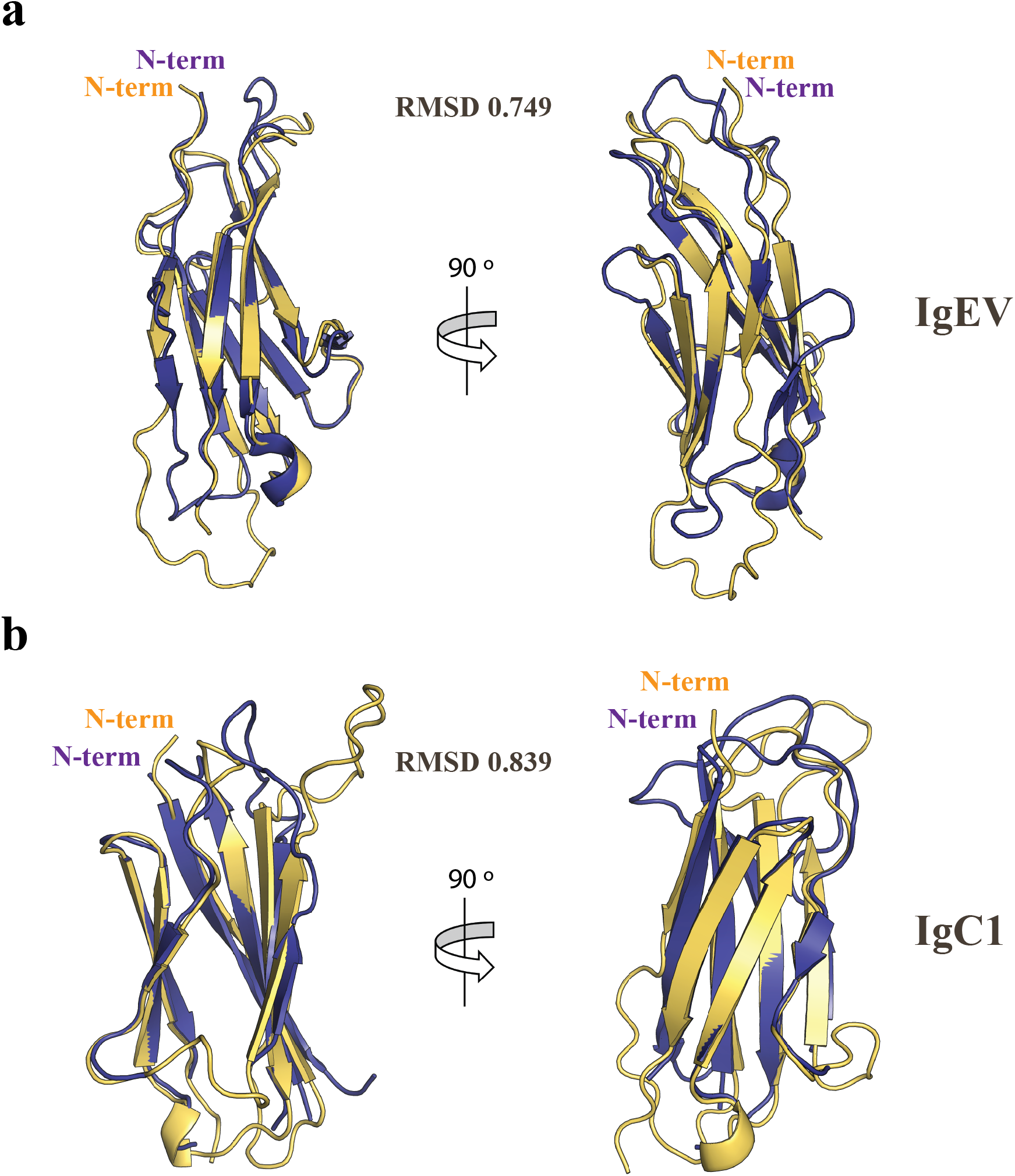
Structural alignment of the Ig individual domains. **A-B**, shown are two views of the experimental structures of the N- (a) or C-terminal (b) Ig-domains and their equivalent AlphaFold models, aligned throught the alpha carbons. The relative N- and C-terminal residues are indicated.

**Fig S3.**
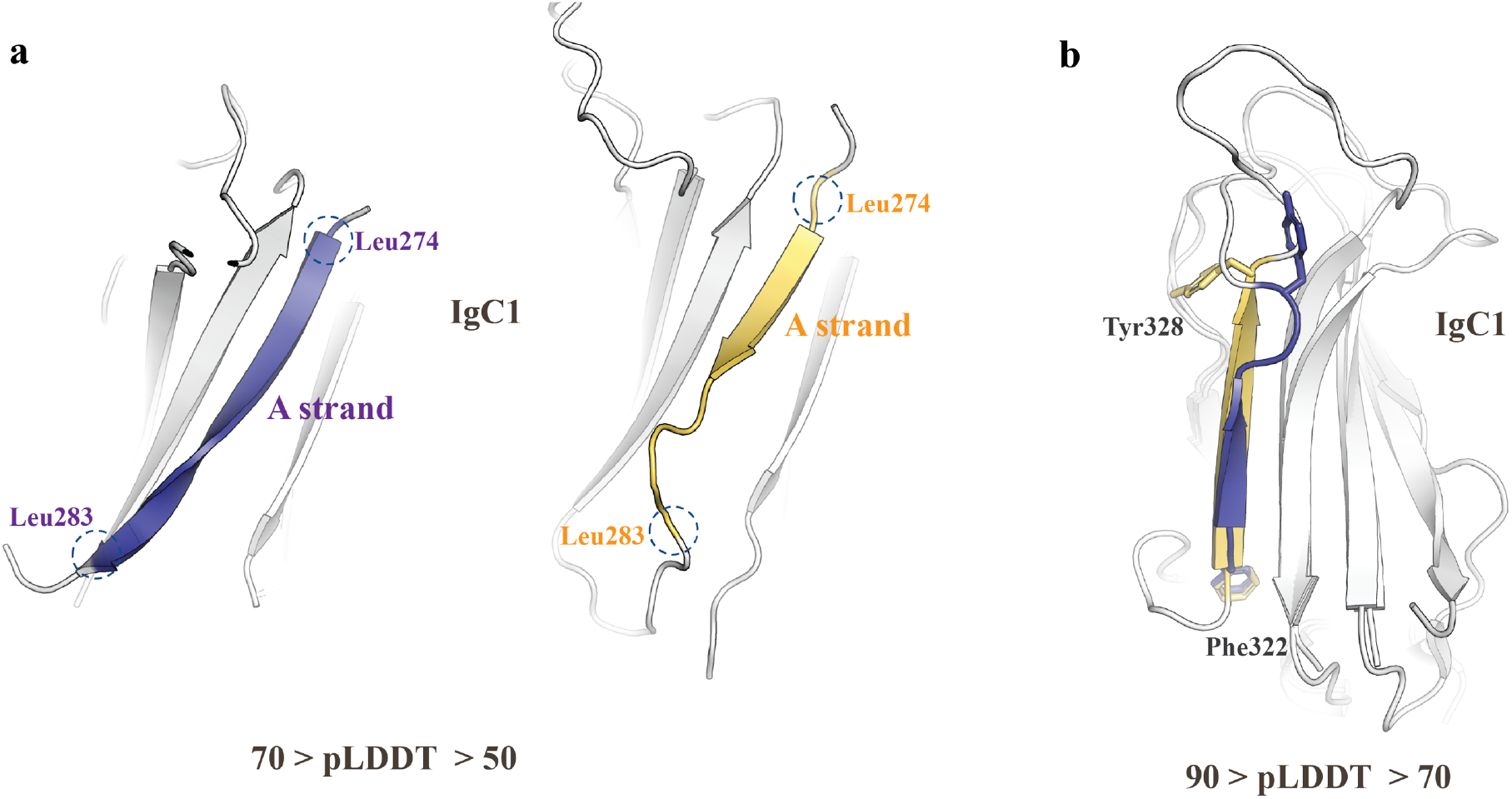
Secondary structure mismatch between the predicted model and the experimental structure. **A**, shown are beta sheet strand motives in the experimental and AlphaFold models corresponding the to the C-terminal IgC1 domain of SAML. The structures are shown light grey colored with the target beta strands in deepblue and yelloworange (AlphaFold prediction) colors. **B**, the C-terminal IgC1 domain of the experimental and AlphaFold structures are superposed through the alpha carbons. The FNITPRY motives are highlighted for visual comparison. Secondary structure assignments were produced in Pymol (version 2.5.2) following the Dictionary of Secondary Structure in Proteins (DSSP) criteria^6^. The model confidence level is included as a pLDDT range.

## Notes

https://www.rcsb.org/structure/8OVQ

### Competing Interest Statement

The authors have declared no competing interest.

